# Enhanced proofreading governs CRISPR-Cas9 targeting accuracy

**DOI:** 10.1101/160036

**Authors:** Janice S. Chen, Yavuz S. Dagdas, Benjamin P. Kleinstiver, Moira M. Welch, Lucas B. Harrington, Samuel H. Sternberg, J. Keith Joung, Ahmet Yildiz, Jennifer A. Doudna

**Affiliations:** Department of Molecular and Cell Biology, University of California, Berkeley, California, 94720, USA; Biophysics Graduate Group, University of California, Berkeley, California 94720, USA; Molecular Pathology Unit, Center for Cancer Research, and Center for Computational and Integrative Biology, Massachusetts General Hospital, Charlestown, Massachusetts 02129, USA; Department of Pathology, Harvard Medical School, Boston, Massachusetts 02115, USA; Department of Chemistry, University of California, Berkeley, California 94720, USA; Department of Physics, University of California, Berkeley, California 94720, USA; Howard Hughes Medical Institute, University of California, Berkeley, California 94720, USA; Physical Biosciences Division, Lawrence Berkeley National Laboratory, Berkeley, California 94720, USA

## Abstract

The RNA-guided CRISPR-Cas9 nuclease from *Streptococcus pyogenes* (SpCas9) has been widely repurposed for genome editing^1-4^. High-fidelity (SpCas9-HF1) and enhanced specificity (eSpCas9(1.1)) variants exhibit substantially reduced off-target cleavage in human cells, but the mechanism of target discrimination and the potential to further improve fidelity were unknown^5-9^. Using single-molecule Förster resonance energy transfer (smFRET) experiments, we show that both SpCas9-HF1 and eSpCas9(1.1) are trapped in an inactive state^10^ when bound to mismatched targets. We find that a non-catalytic domain within Cas9, REC3, recognizes target mismatches and governs the HNH nuclease to regulate overall catalytic competence. Exploiting this observation, we identified residues within REC3 involved in mismatch sensing and designed a new hyper-accurate Cas9 variant (HypaCas9) that retains robust on-target activity in human cells. These results offer a more comprehensive model to rationalize and modify the balance between target recognition and nuclease activation for precision genome editing.

---

Efforts to minimize off-target cleavage by CRISPR-Cas9 have motivated the development of SpCas9-HF1 and eSpCas9(1.1) variants that contain amino acid substitutions predicted to weaken the energetics of target site recognition and cleavage^8,9^ **(Figure 1a)**. Biochemically, we found that these Cas9 variants cleaved the on-target DNA with rates similar to that of wild-type (WT) SpCas9, whereas their cleavage activity was significantly reduced on substrates bearing mismatches **(Extended Data Figure 1a, 2a)**. To test the hypothesis that SpCas9 with its single-guide RNA (sgRNA) might exhibit a greater affinity for its target than is required for effective recognition^9,11^, we measured DNA binding affinity and cleavage of SpCas9-HF1 and eSpCas9(1.1) variants. Contrary to a potential hypothesis that mutating these charged residues to alanine weakens target binding^11^, the affinities of these variants for on-target and PAM-distal mismatched substrates were similar to WT SpCas9 **(Figure 1b, Extended Data Figure 1a, 2b)**, indicating that cleavage specificity is improved through a mechanism distinct from a simple reduction of target binding affinity^11^.

**Figure 1.**
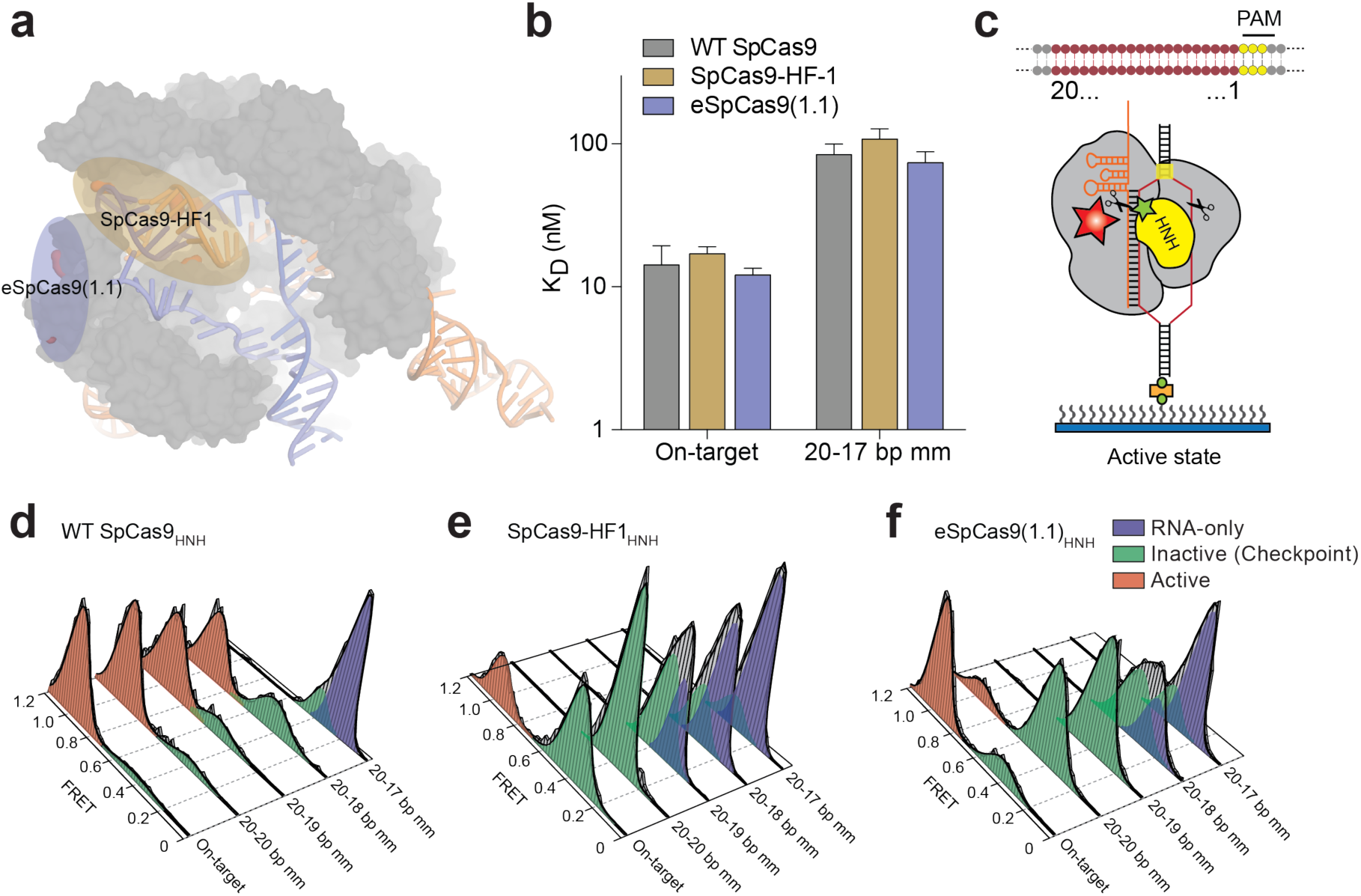
High-fidelity Cas9 variants enhance cleavage specificity through HNH conformational control. **a,** Locations of amino acid alterations present in existing high-fidelity SpCas9 variants mapped onto the dsDNA-bound SpCas9 crystal structure (5F9R), with the HNH domain omitted for clarity. **b,** Dissociation constants comparing WT SpCas9, SpCas9-HF1 and eSpCas9(1.1) with perfect and a 20-17 bp mismatched target. Error bars, s.d.; *n* = 3. **c,** Cartoon of DNA-immobilized SpCas9 complexes for smFRET experiments with DNA target numbering scheme. **d–f,** smFRET histograms measuring HNH conformational activation with **d,** WT SpCas9_HNH_, **e,** SpCas9-HF1_HNH_ and **f,** eSpCas9(1.1)_HNH_ bound to perfect and PAM-distal mismatched targets. Black curves represent a fit to multiple Gaussian peaks.

The HNH nuclease domain of SpCas9 undergoes a substantial conformational rearrangement upon target binding^12-15^, which activates the RuvC nuclease for concerted cleavage of both strands of the DNA^12,16^. We have previously shown that the HNH domain stably docks in its active state with an on-target substrate but becomes loosely trapped in an catalytically-inactive conformational checkpoint when bound to mismatched targets^10^, suggesting that SpCas9-HF1 and eSpCas9(1.1) variants may employ a more stringent checkpoint to promote off-target discrimination. To test this possibility, we labeled catalytically active WT SpCas9 (SpCas9_HNH_), SpCas9-HF1 (SpCas9-HF1_HNH_) and eSpCas9(1.1) (eSpCas9(1.1)_HNH_) with Cy3/Cy5 FRET pairs at positions S355C and S867C to measure HNH conformational states **(Figure 1c–f, Extended Data Figure 1c–e)**^12^. Whereas SpCas9_HNH_ stably populated the active state with both on-target and mismatched substrates in steady-state smFRET histograms **(Figure 1d)**, only ∼32% of SpCas9-HF1_HNH_ molecules occupied the HNH active state (E_FRET_ = 0.97) with an on-target substrate, with the remaining ∼68% trapped in the inactive intermediate state (E_FRET_ = 0.45) **(Figure 1e)**. However, when SpCas9-HF1_HNH_ was bound to a substrate with just a single nucleotide mismatch at the PAM-distal end (20-20 bp mm), stable docking of the HNH nuclease was entirely ablated **(Figure 1e)**. In addition, eSpCas9(1.1)_HNH_ and other high fidelity variants^8,9^ reduced the HNH active state in the presence of mismatches **(Figure 1f, Extended Data Figure 2c–d)**. We therefore propose that high fidelity variants of Cas9 reduce off-target cleavage by raising the threshold for HNH conformational activation when bound to DNA substrates.

**Figure 2.**
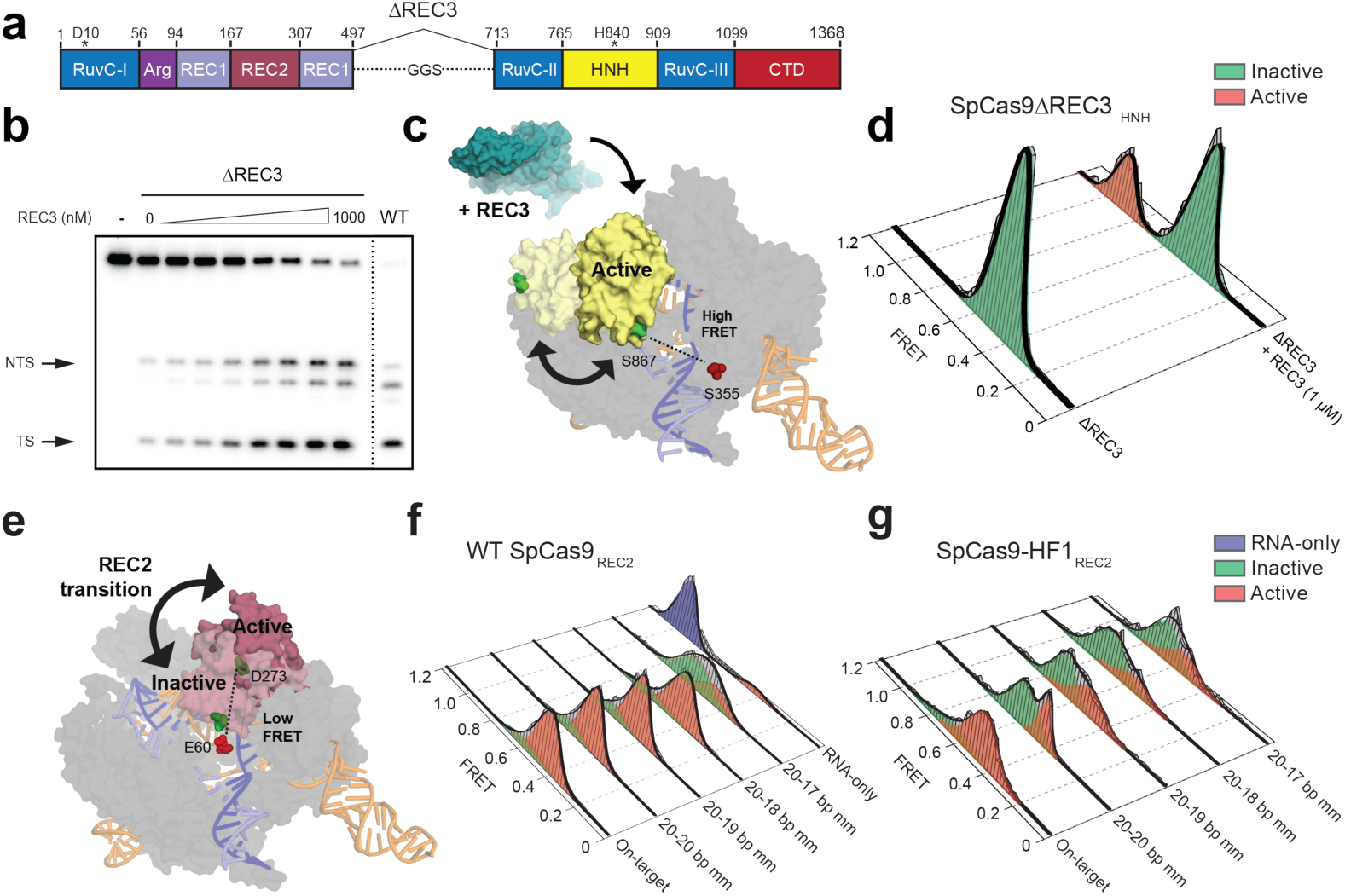
The alpha-helical lobe regulates HNH domain activation. **a,** Domain organization of SpCas9ΔREC3. **b,** Perfect target DNA cleavage assay using SpCas9ΔREC3 with increasing concentrations of REC3 domain supplied in *trans*, resolved by denaturing PAGE. **c,** Schematic of SpCas9ΔREC3_HNH_ with FRET dyes at positions S355C and S867C, with the REC3 domain added *in trans*. Inactive to active structures represent HNH in the sgRNA-bound (PDB ID: 4ZT0) to dsDNA-bound (PDB ID: 5F9R) forms, respectively. **d,** smFRET histograms measuring HNH conformational states with SpCas9ΔREC3_HNH_ in the absence and presence of the REC3 domain. **e,** Schematic of SpCas9_REC2_ with FRET dyes at positions E60C and D273C, with HNH domain omitted for clarity. Inactive to active structures represent REC2 in the sgRNA-bound (PDB ID: 4ZT0) to dsDNA-bound (PDB ID: 5F9R) forms, respectively. **f–g,** smFRET histograms measuring REC2 conformational states with **f,** WT SpCas9_REC2_ and **g,** SpCas9-HF1_REC2_ bound to perfect and PAM-distal mismatched targets. For panels **d, f** and **g,** black curves represent a fit to multiple Gaussian peaks.

Since the HNH domain does not directly contact nucleic acids at the PAM-distal end^13,17-19^, it is likely that a separate domain of Cas9 senses mismatches to govern HNH domain mobility. Structural studies suggest that a domain within the Cas9 recognition (REC) lobe (REC3) interacts with the RNA/DNA heteroduplex and undergoes conformational changes upon target binding **(Extended Data Figure 3)**^13,14,17-19^. Because the function of this non-catalytic domain was previously unknown, we labeled SpCas9 with Cy3/Cy5 dyes at positions S701C and S960C (SpCas9_REC3_) and observed that the conformational states of REC3 become more heterogeneous as PAM-distal mismatches increase **(Extended Data Figure 4a-c)**. To determine whether PAM-distal sensing precedes HNH activation, we deleted REC3 from WT Cas9 (SpCas9ΔREC3) **(Figure 2a)**. Deletion of REC3 decreased the cleavage rate by ∼1000-fold compared to WT Cas9, despite retaining near-WT binding affinity with a perfect target **(Extended Data Figure 4d–f)**. Unexpectedly, *in vitro* complementation of REC3 domain *in trans* rescued the on-target cleavage rate by ∼100-fold in a concentration-dependent manner, but had no effect on cleavage with a PAM-distal mismatched target **(Figure 2b, Extended Data Figure 4e)**. Furthermore, smFRET experiments revealed that the HNH domain in SpCas9ΔREC3 (SpCas9Δ REC3_HNH_) occupied the active state only when REC3 was supplemented *in trans* **(Figure 2c–d, Extended Data Figure 4f)**. We therefore propose that REC3 acts as an allosteric effector that recognizes RNA/DNA heteroduplex to allow for HNH nuclease activation.

We next considered allosteric interactions that could couple the discontinuous REC3 and HNH domains. Structural studies suggested that REC2 occludes the HNH domain from the scissile phosphate in the sgRNA-bound state^19^, and undergoes a large outward rotation upon binding to double-stranded DNA (dsDNA)^13,14^ **(Figure 2e)**. To test whether the REC2 domain regulates access of HNH to the target strand scissile phosphate, we labeled SpCas9 with Cy3/Cy5 dyes at positions E60C and D273C (SpCas9_REC2_) in order to detect REC2 conformational changes **(Extended Data Figure 1b–c)**. We observed reciprocal changes in bulk FRET values ((ratio)_A_)^20^ between SpCas9_HNH_ and SpCas9_REC2_ across multiple DNA substrates **(Extended Data Figure 4g)**, which suggest that the REC2 and HNH domains are tightly coupled to ensure catalytic competence. smFRET experiments further confirmed a large opening of REC2 during the transition from the sgRNA-bound state (E_FRET_ = 0.96) to the target-bound state (E_FRET_ = 0.43) **(Figure 2e–f)**. In contrast to WT SpCas9_REC2_, SpCas9-HF1_REC2_ occupies an intermediate state (E_FRET_ = 0.63) when bound to a target with just a single PAM-distal mismatch **(Figure 2f–g)**. Together with the observation that the HNH domain of SpCas9-HF1 does not occupy the active state in the presence of PAM-distal mismatches, these experiments suggest that REC2 sterically occludes and traps the HNH nuclease domain in the conformational checkpoint when SpCas9 is bound to off-target substrates.

Next, we investigated if this conformational proofreading mechanism could be rationally exploited to design a suite of novel hyper-accurate Cas9 variants. We identified five clusters of residues containing conserved amino acids within 5 Å of the RNA/DNA interface, four of which are located within REC3 and one in the HNH-RuvC Linker 2 (L2) **(Figure 3a, Extended Data Figure 5)**. Alone or in combination with Q926A, a substitution within L2 that confers specificity^9^, we generated alanine substitutions for each residue within five different clusters of amino acids (Clusters 1-5 ± Q926A) **(Figure 3a)**. We tested whether these cluster mutations affected off-target discrimination and equilibrium binding *in vitro*, and found that Cluster 1 alone and Cluster 2 + Q926A exhibited the greatest suppression of off-target cleavage while retaining target binding affinities comparable to WT **(Extended Data Figure 6)**. We next screened all cluster variants in human cells using an enhanced GFP (*EGFP*) disruption assay^5^. On-target activity for Cluster 1 was comparable to that of SpCas9-HF1 or eSpCas9(1.1), whereas Cluster 2 variants displayed generally lower activity **(Figure 3b, Extended Data Figure 7a)**. Furthermore, Cluster 1 retained high on-target activity (> 70% of WT) at 19/24 endogenous gene sites tested, compared to 18/24 for SpCas9-HF1 and 23/24 for eSpCas9(1.1) **(Figure 3c, Extended Data Figure 8a)**.

**Figure 3.**
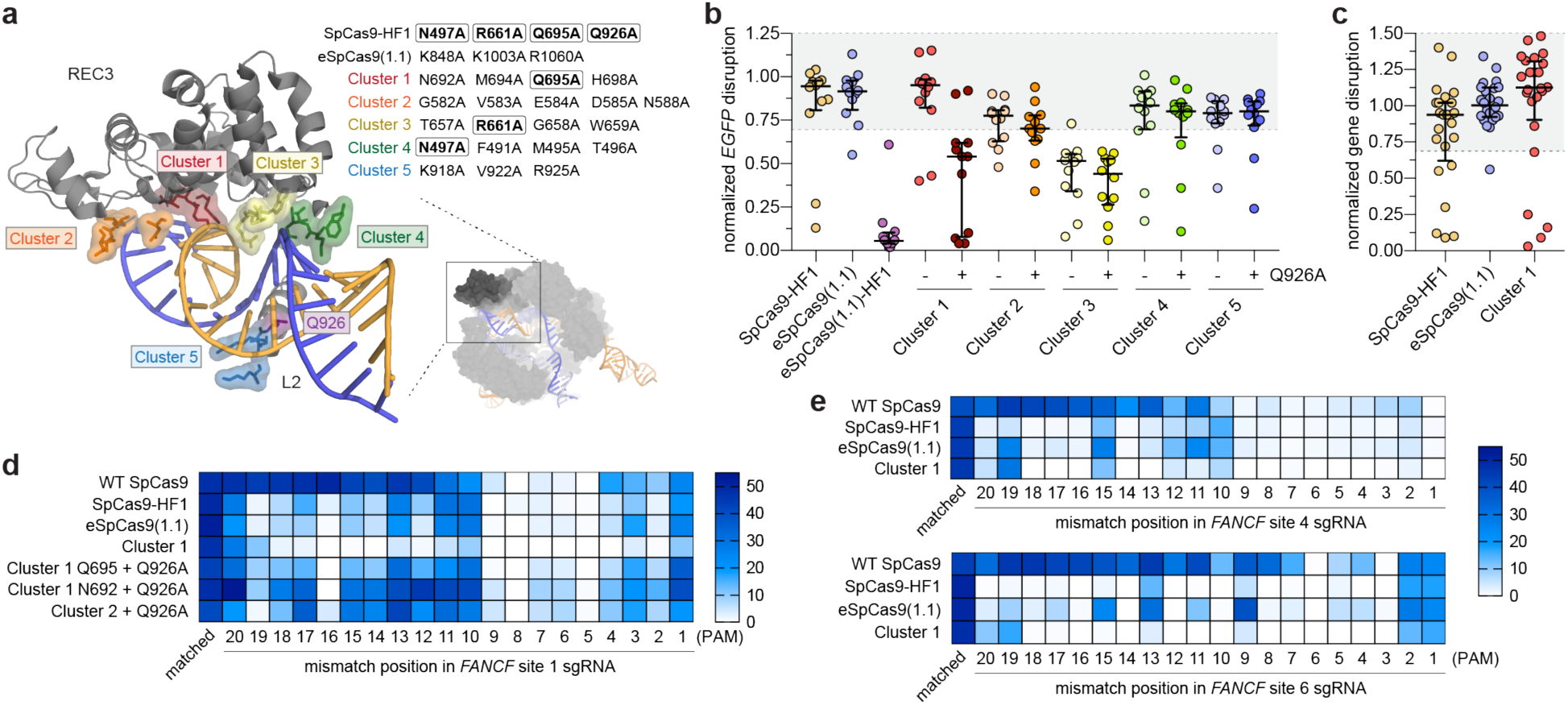
Targeted mutagenesis within the REC3 domain reveals a SpCas9 variant with hyper-accurate behavior in human cells. **a,** Zoomed image of the REC3 domain and Linker 2 (L2) with Cluster variants indicated. Boxed residues indicate amino acids also present in SpCas9-HF1. **b,** WT-normalized activity of SpCas9-HF1, eSpCas9(1.1) and Cluster variants, using sgRNAs targeting 12 different sites within *EGFP*. **c,** WT-normalized endogenous gene disruption activity measured by T7 endonuclease 1 (T7E1) assay across 24 sites. **d,** Activities of WT and high-fidelity Cas9 variants when programmed with singly mismatched sgRNAs against *FANCF* site 1 as measured by T7E1 assay. **e,** Activities of WT SpCas9, SpCas9-HF1, eSpCas9(1.1) and Cluster 1 when programmed with singly mismatched sgRNAs against *FANCF* site 4 and *FANCF* site 6. For panels **b** and **c**, error bars represent median and interquartile ranges, and n ≥ 3; the interval with > 70% of wild-type activity is highlighted in light grey.

We then focused on the specific contributions of mutations within Cluster 1 by restoring each individual mutated residue to its wild-type identity, along with the Q926A mutation, and testing the resulting variants for on-target editing efficiency in human cells. On-target activity was significantly compromised when N692A/Q695A/Q926A mutations occurred together, but restoring either N692 (Cluster 1 N692 + Q926A) or Q695 (Cluster 1 Q695 + Q926A) alone led to robust on-target efficiency comparable to Cluster 1, signifying differential contributions from these mutations to activity and specificity (**Extended Data Figure 7b-c, 8a-b**). Using sgRNAs with single mismatches against the endogenous human gene target *FANCF* site 1, we found that Cluster 1 exhibited even greater specificity than both SpCas9-HF1 and eSpCas9(1.1) in the middle and PAM proximal regions of the spacer, suggesting that mutating N692A and Q695A together may induce specificity in parts of the spacer sequence that were previously susceptible to off-target cleavage by high-fidelity Cas9 variants^9^ **(Figure 3d, Extended Data Figure 8c)**. Additional single mismatch tolerance assays on *FANCF* sites 4 and 6 further corroborated the superior accuracy of Cluster 1 (N692A/M694A/Q695A/H698A, referred to as HypaCas9) against mismatches at positions 1 through 18; however, single mismatches along *FANCF* site 2 were still tolerated across all SpCas9 variants tested **(Figure 3e, Extended Data Figure 8d, e)**.

To biochemically validate cleavage specificity in the middle region of the spacer with HypaCas9, we measured cleavage rates against the *FANCF* site 1 sequence with or without internal mismatches at positions 10-12 of the spacer. Although HypaCas9 retained on-target activity comparable to WT and SpCas9-HF1 in human cells, its *in vitro* cleavage rate was slightly reduced for the one target site examined **(Figure 4a)**. However, the cleavage rate with internally mismatched substrates was considerably slower compared to WT and SpCas9-HF1 **(Figure 4a)**. This activity may be explained by the altered threshold of HNH domain activation; whereas stable HNH docking was observed by SpCas9_HNH_ and SpCas9-HF1_HNH_ with both the *FANCF* site 1 on-target and mismatched substrate at the 12^th^ position, this HNH active state by HypaCas9 (HypaCas9_HNH_) was diminished with the on-target. Nevertheless, HNH docking was nearly abolished when HypaCas9_HNH_ was bound to a substrate with a single mismatch **(Figure 4b)**.

**Figure 4.**
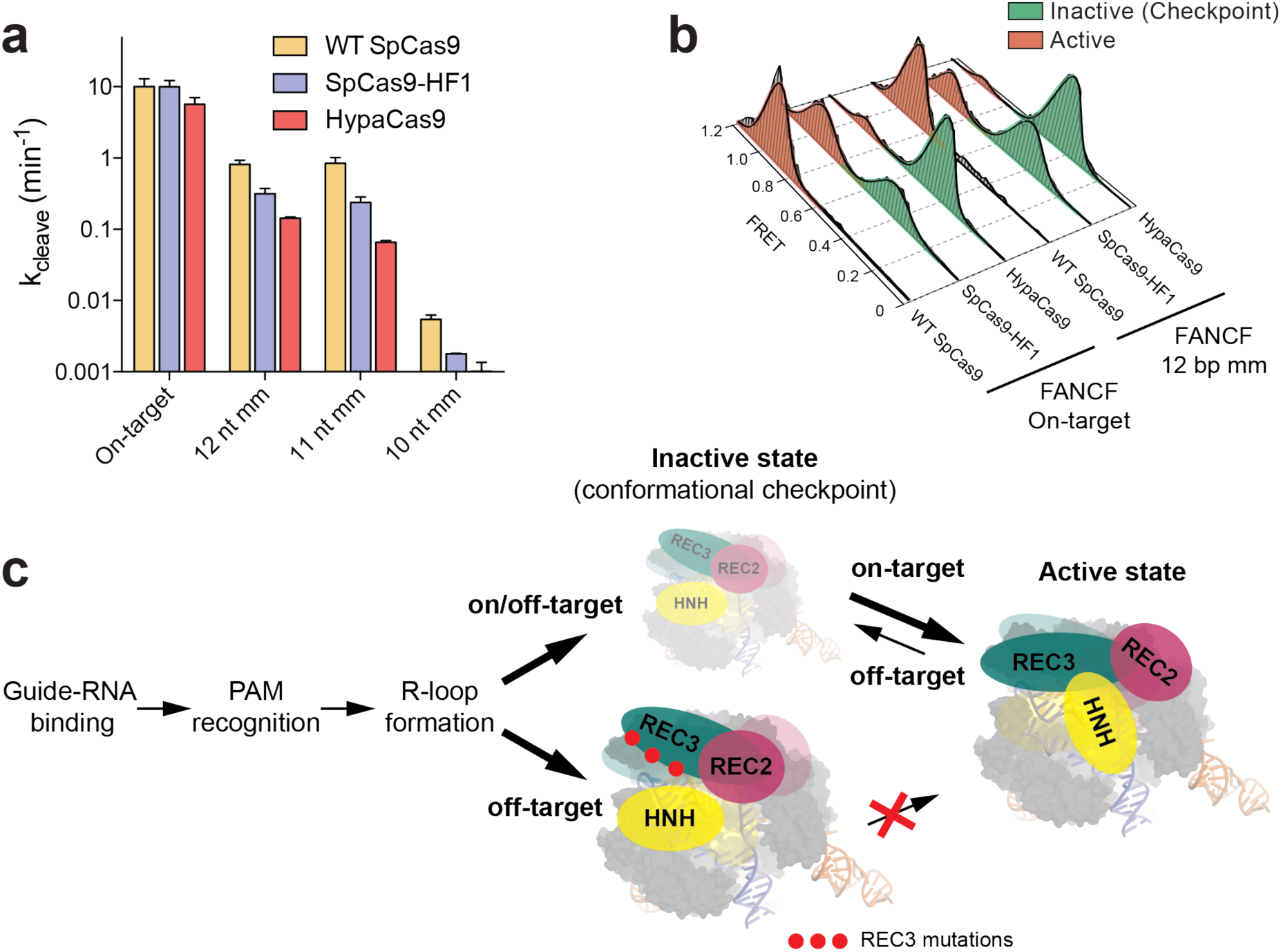
Mutating residues involved in proofreading increases the threshold for conformational activation to ensure targeting accuracy. **a,** DNA cleavage kinetics of SpCas9 variants with the *FANCF* site 1 on-target and internally mismatched substrates. Error bars, s.d.; *n* = 3. **b,** smFRET histograms measuring HNH conformational states for indicated SpCas9 variants with a *FANCF* site 1 on-target and mismatched substrate at the 12^th^ position; black curves represent a fit to multiple Gaussian peaks. **c,** Model for alpha-helical lobe sensing and regulation of the RNA/DNA heteroduplex for HNH activation and cleavage.

Our findings provide direct evidence to support previous speculation that Cas9 relies on PAM-distal protospacer sensing to enable accurate targeting^21,22^. In particular, we define REC3 as an allosteric regulator of global Cas9 conformational changes to activate the nuclease domains, whose conformational threshold can be tuned for high-fidelity cleavage. Mutation of residues within REC3 that are involved in nucleic acid recognition, such as those mutated in HypaCas9 or SpCas9-HF1, prevents transitions by the REC lobe, which more stringently traps the HNH domain in the conformational checkpoint in the presence of mismatches **(Figure 4c, Extended Data Figure 9)**. Curiously, nearly all of the amino acids within the cluster variants were strongly conserved **(Extended Data Figure 5)**, suggesting that these residues may also be involved in protospacer sensing and HNH nuclease activation across Cas9 orthologues. Furthermore, this observation may address how nature apparently has not selected for a highly precise Cas9 protein, whose native balance between mismatch tolerance and specificity may be optimized for host immunity. Our study therefore delineates a general strategy for improving Cas9 specificity by tuning conformational activation and offers innovative opportunities for rational design of hyper-accurate Cas9 variants that do not compromise efficiency.

## METHODS

### Protein purification and dye labeling

*S. pyogenes* Cas9 and truncation derivatives were cloned into a custom pET-based expression vector containing an N-terminal His_6_-tag, maltose-binding protein (MBP) and TEV protease cleavage site. Point mutations were introduced by Gibson assembly or around-the-horn PCR and verified by DNA sequencing. Proteins were purified as described^23^, with the following modifications: after Ni-NTA affinity purification and overnight TEV cleavage at 4°C, proteins were purified over an MBPTrap HP column connected to a HiTrap Heparin HP column for cation exchange chromatography. The final gel filtration step (Superdex 200) was carried out in elution buffer containing 20 mM Tris-HCl pH 7.5, 200 mM NaCl, 5% glycerol (v/v) and 1 mM TCEP. For FRET experiments, dye-labeled Cas9 samples were prepared as described^12^. A list of all protein variants and truncations are listed in **Supplementary Table 1**.

### Nucleic acid preparation

sgRNA templates were PCR amplified from a pUC19 vector containing a T7 promoter, 20 nt target sequence and optimized sgRNA scaffold. The amplified PCR product was extracted with phenol:chloroform:isoamylalcohol and served as the DNA template for sgRNA transcription reactions, which were performed as described^24^. DNA oligonucleotides and 5′end biotinylated DNAs (Supplementary Table 2) were synthesized commercially (Integrated DNA Technologies), and DNA duplexes were prepared and purified by native PAGE as described^23^.

### DNA cleavage and binding assays

DNA duplex substrates were 5′-[^32^P]-radiolabeled on both strands. For cleavage experiments, Cas9 and sgRNA were pre-incubated at room temperature for at least 10 min in 1X binding buffer (20 mM Tris-HCl pH 7.5, 100 mM KCl, 5 mM MgCl_2_, 1 mM DTT, 5% glycerol, 50 µg ml^-1^ heparin) before initiating the cleavage reaction by addition of DNA duplexes. For REC3 *in vitro* complementation experiments, SpCas9ΔREC3 and sgRNA were pre-incubated with 10-fold molar excess of REC3 for at least 10 minutes at room temperature before addition of radiolabeled substrate. DNA cleavage experiments were performed and analyzed as previously described^12^. DNA binding assays were conducted in 1X binding buffer without MgCl_2_ + 1 mM EDTA at room temperature for 2 hours. DNA-bound complexes were resolved on 8% native PAGE (0.5X TBE + 1 mM EDTA, without MgCl_2_) at 4°C, as previously described^10^. Experiments were replicated at least three times, and presented gels are representative results.

### Bulk FRET experiments

All bulk FRET assays were performed at room temperature in 1× binding buffer, containing 50 nM SpCas9_HNH_ (C80S/S355C/C574S/S867C labeled with Cy3/Cy5), SpCas9ΔREC3_HNH_(M1–N497,GGS,V713–D1368 + C80S/S355C/C574S/S867C) or SpCas9_REC2_ (E60C/C80S/D273C/C574S labeled with Cy3/Cy5) with 200 nM sgRNA and DNA substrate where indicated. Fluorescence measurements were collected and analyzed as described^12^. For REC3 *in vitro* complementation FRET experiments, SpCas9ΔREC3_HNH_ and sgRNA were pre-incubated with 10-fold molar excess of REC3 for at least 10 minutes at room temperature before measuring bulk fluorescence.

### Sample preparation for smFRET assay

99% PEG and 1% biotinylated-PEG coated quartz slides were received from MicroSurfaces, Inc. Sample preparation was performed as previously described^10^. To immobilize SpCas9 on its DNA substrate, 2.5nM biotinylated-DNA substrate introduced and incubated in sample chamber for 5 min. Excess DNA was washed with 1× binding buffer. SpCas9-sgRNA complexes were prepared by mixing 50 nM Cas9 and 50nM sgRNA in 1× binding buffer and incubated for 10 min at room temperature. SpCas9-sgRNA was diluted to 100 pM, introduced to sample chamber and incubated for 10 min. Before data acquisition, 20 µL imaging buffer (1 mg ml^-1^ glucose oxidase, 0.04 mg ml^-1^ catalase, 0.8% dextrose (w/v) and 2 mM Trolox in 1× binding buffer) was flown into chamber. The REC3 *in vitro* complementation assay was performed similar to steady-state FRET experiments: 2.5nM biotinylated-DNA substrate (on-target) was immobilized on surface, and excess DNA was washed with 1× binding buffer. SpCas9-sgRNA complexes were prepared by mixing 50 nM SpCas9ΔREC3 and 50nM sgRNA in 1× binding buffer and incubated for 10 min at room temperature. SpCas9-sgRNA was diluted to 100 pM, introduced to the sample chamber and incubated for 10 min. Before data acquisition, 20 µL imaging buffer was flown into chamber. After data acquisition, the sample chamber was washed with 1× binding buffer. 20 µL imaging buffer supplemented with 1µM REC3 was flown into sample chamber and incubated for 10min. After incubation, data for REC3 complementation was collected.

### Microscopy and data analysis

A prism-type TIRF microscope was setup using a Nikon Ti-E Eclipse inverted fluorescent microscope equipped with a 60× 1.20 N.A. Plan Apo water objective and the perfect focusing system (Nikon). A 532-nm solid state laser (Coherent Compass) and a 633-nm HeNe laser (JDSU) were used for Cy3 and Cy5 excitation, respectively. Cy3 and Cy5 fluorescence were split into two channels using an Optosplit II image splitter (Cairn Instruments) and imaged separately on the same electron-multiplied charged-coupled device (EM-CCD) camera (512×512 pixels, Andor Ixon EM^+^). Effective pixel size of the camera was set to 267 nm after magnification. Movies for steady-state FRET measurements were acquired at 10 Hz under 0.3 kW cm^-2^ 532-nm excitation. Data analysis was performed as described previously^10^. Briefly, two fluorescent channels were registered with each other using fiducial markers (20 nm diameter Nile Red Beads, Life Technologies) to determine the Cy3/Cy5 FRET pairs. Cy3/Cy5 pairs that photobleached in one step and showed anti-correlated signal changes were used to build histograms. FRET values were corrected for donor leakage and the histograms were normalized to determine the percentage of distinct FRET populations.

### Human cell culture and transfection

Descriptions of nuclease and guide RNA plasmids used for human cell culture are available in **Supplementary Table 1 and 2**. Nuclease variants were generated by isothermal assembly into JDS246 (Addgene #43861)^5^, and guide RNAs were cloned into BsmBI digested BPK1520 (Addgene #65777)^25^. Both U2OS cells (a gift from Toni Cathomen, Freiburg) and U2OS-EGFP cells (encoding a single integrated copy of a pCMV-EGFP–PEST cassette)^26^ were cultured at 37°°C with 5% CO_2_ in advanced DMEM containing 10% heat-inactivated fetal bovine serum, 2 mM GlutaMax, penicillin/streptomycin, and 400 µg ml^-1^ Geneticin (for U2OS-EGFP cells only). Cell culture reagents were purchased from Thermo Fisher Scientific, cell line identities were validated by STR profiling (ATCC) and deep-sequencing, and cell culture supernatant was tested bi-weekly for mycoplasma. Transfections were performed using a Lonza 4-D Nucleofector with the SE Kit and the DN-100 program on ∼200k cells with 750 ng of nuclease and 250 ng of guide RNA plasmids.

### Human cell EGFP disruption assay

*EGFP* disruption experiments were performed as previously described^5,26^. Briefly, transfected cells were analyzed ∼52 hours post-transfection for loss of EGFP fluorescence using a Fortessa flow cytometer (BD Biosciences). Background loss was determined by gating a negative control transfection (containing nuclease and empty guide RNA plasmid) at ∼2.5% for all experiments.

### T7 endonuclease I assay

Roughly 72 hours post-transfection, genomic DNA was extracted from U2OS cells using the Agencourt DNAdvance Genomic DNA Isolation Kit (Beckman Coulter Genomics), and T7 endonuclease I assays were performed as previously described^26^. Briefly, 600**–**800 nt amplicons surrounding on-target sites were amplified from ∼100 ng of genomic DNA using Phusion Hot-Start Flex DNA Polymerase (New England Biolabs) using the primers listed in Supplementary Table 2. PCR products were visualized (using a QIAxcel capillary electrophoresis instrument, Qiagen), and purified (Agencourt Ampure XP cleanup, Beckman Coulter Genomics), Denaturation and annealing of ∼200 ng of the PCR product was followed by digestion with T7 endonuclease I (New England Biolabs). Digestion products were purified (Ampure) and quantified (QIAxcel) to approximate the mutagenesis frequencies induced by Cas9-sgRNA complexes.

## ACKNOWLEDGEMENTS

We thank Addison V. Wright, Stephen N. Floor, Joshua C. Cofsky, David Burstein, Christof Fellman, Benjamin L. Oakes and Orestes Mavrothalassitis for discussions and critical reading of the manuscript, and Michelle S. Prew for technical assistance. J.S.C. acknowledges support from the National Science Foundation Graduate Research Fellowship program, and B.P.K. from Banting (Natural Sciences and Engineering Research Council of Canada) and Charles A. King Trust Postdoctoral Fellowships. J.A.D. is an Investigator of the Howard Hughes Medical Institute. This work has been supported by NIH (GM094522 and GM118773 (A.Y.), R35 GM118158 (J.K.J.)), NSF (MCB-1617028 (A.Y.) and MCB-1244557 (J.A.D.)), and the Desmond and Ann Heathwood MGH Research Scholar Award (J.K.J.).

## AUTHOR CONTRIBUTIONS

J.S.C., Y.S.D. and B.P.K. conceived of and designed experiments with input from L.B.H., S.H.S, J.K.J., A.Y. and J.A.D. J.S.C. performed protein expression, labeling and biochemical experiments. Y.S.D. performed single-molecule fluorescence assays and data analysis. B.P.K. and M.M.W. performed human cell-based experiments. J.S.C., Y.S.D., B.P.K., J.K.J., A.Y. and J.A.D. wrote the manuscript.

## COMPETING FINANCIAL INTERESTS

J.K.J. has financial interests in Beacon Genomics, Beam Therapeutics, Editas Medicine, Pairwise Plants, Poseida Therapeutics, and Transposagen Biopharmaceuticals. J.K.J.’s interests were reviewed and are managed by Massachusetts General Hospital and Partners HealthCare in accordance with their conflict of interest policies. J.A.D. is a co-founder of Caribou Biosciences, Editas Medicine, and Intellia Therapeutics; a scientific advisor to Caribou, Intellia, eFFECTOR Therapeutics and Driver; and executive director of the Innovative Genomics Institute at UC Berkeley and UCSF. S.H.S. is an employee of Caribou Biosciences, Inc. and an inventor on patent applications related related to CRISPR-Cas systems and uses thereof. J.S.C, Y.S.D., B.P.K, L.B.H., S.H.S., A.Y., J.K.J. and J.A.D. are inventors on patents for CRISPR technologies.

**Extended Data Figure 1.**
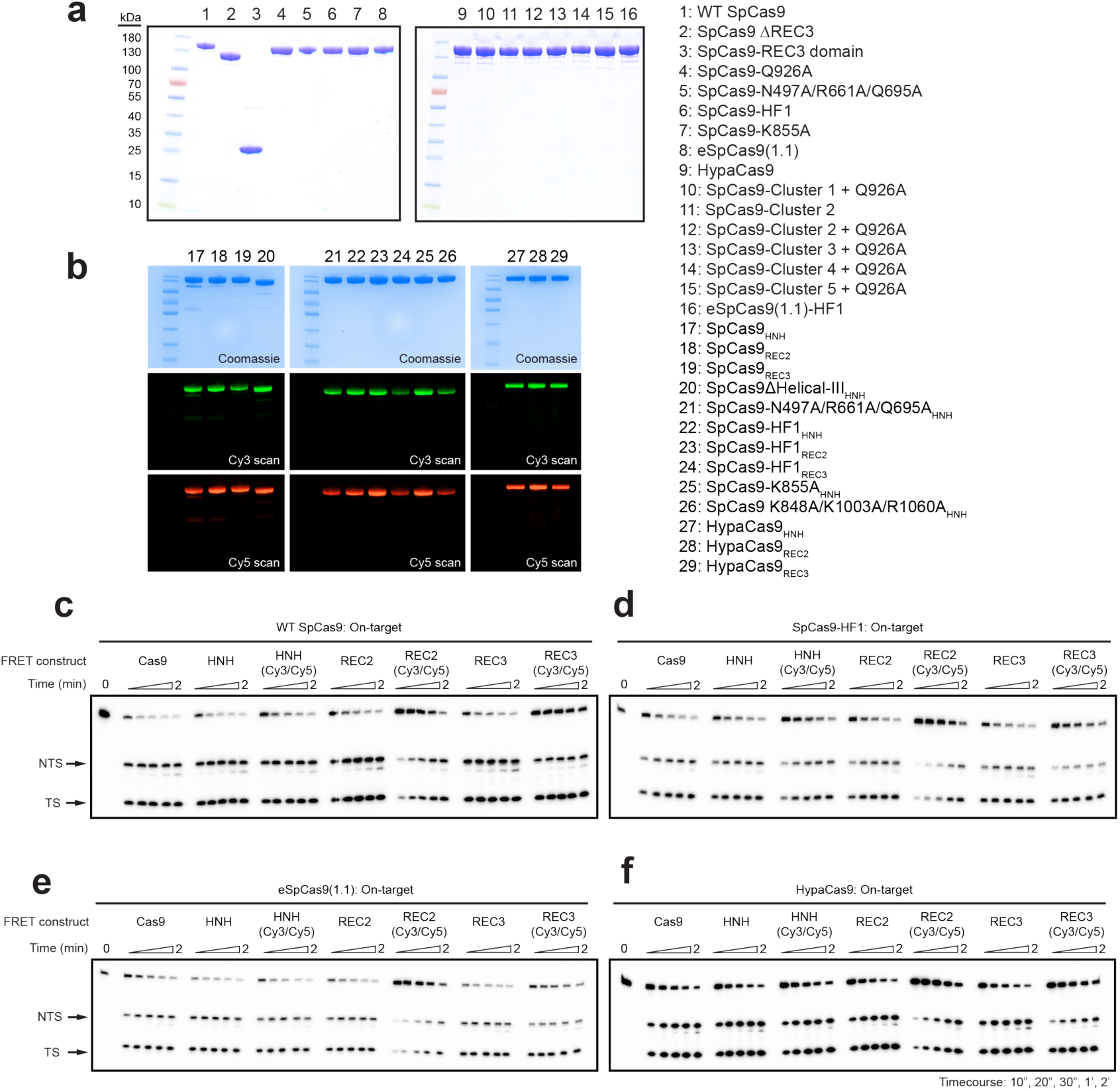
Dually-labeled SpCas9 variants are fully functional for DNA cleavage. **a,** Sodium dodecyl sulphate–polyacrylamide gel electrophoresis (SDS–PAGE) analysis of unlabeled Cas9 variants. **b,** SDS-PAGE analysis of Cy3/Cy5-labeled Cas9 variants. The gel was scanned for Cy3/Cy5 fluorescence (middle, bottom) before staining with Coomassie blue (top). **c–f,** DNA cleavage time courses of Cas9 FRET constructs and their dually-labeled counterparts for **c,** WT SpCas9, **d,** SpCas9-HF1, **e,** eSpCas9(1.1) and **f**, HypaCas9.

**Extended Data Figure 2.**
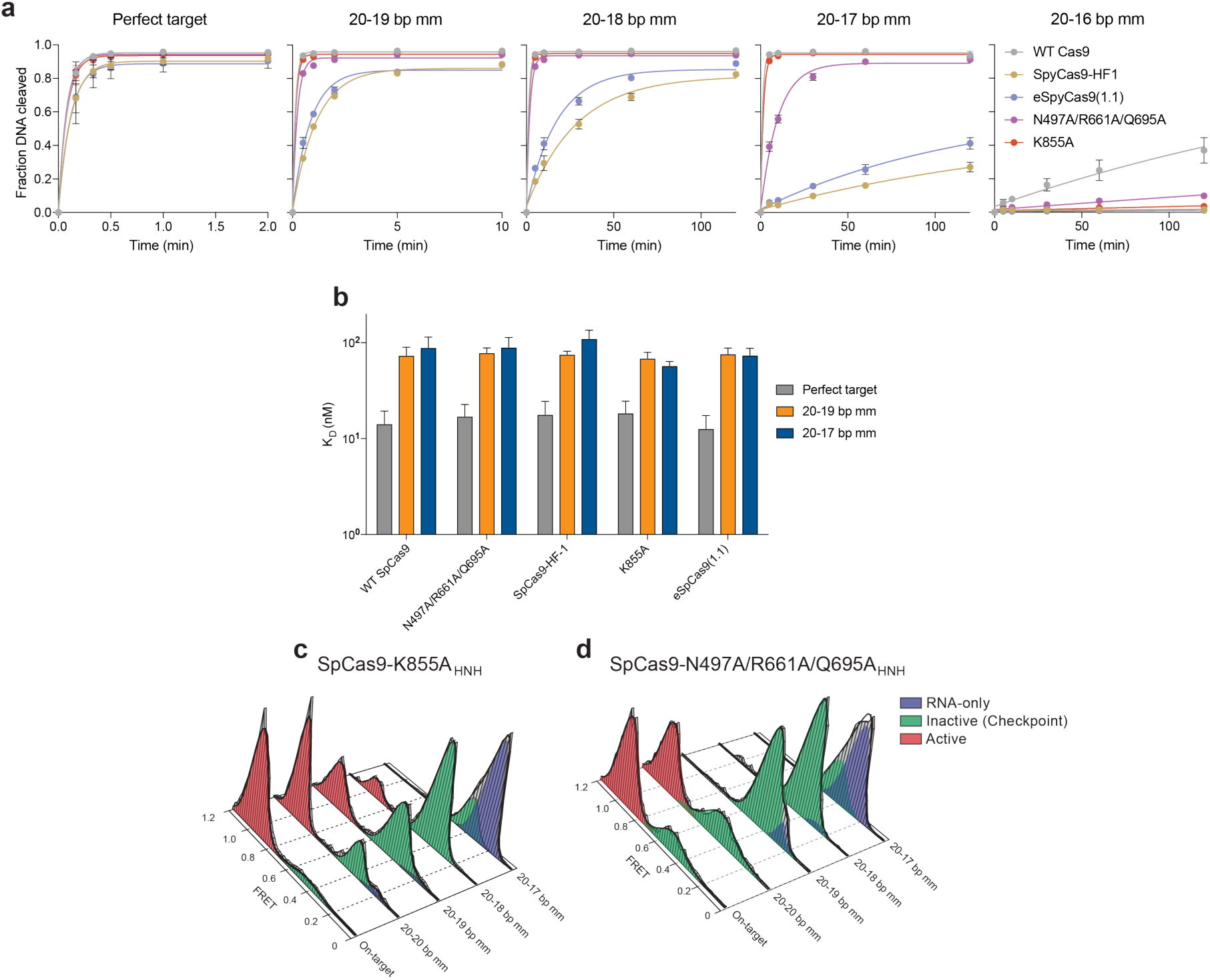
HNH domain in eSpCas9 variants still populate the docked state in the presence of PAM-distal mismatches. **a,** Quantification of DNA cleavage time courses comparing WT SpCas9, SpCas9-HF and eSpCas9(1.1) variants with perfect and PAM-distal mismatched targets. **b,** Dissociation constants comparing WT SpCas9, SpCas9-HF and eSpCas9(1.1) variants with perfect and PAM-distal mismatched targets, as measured by electrophoretic mobility shift assays. Error bars in **a** and **b**, s.d.; *n* = 3. **c–d,** smFRET histograms for **c,** SpCas9-K855A and **d,** SpCas9-N497A/R661A/Q695A. For panels **c** and **d**, black curves represent a fit to multiple Gaussian peaks.

**Extended Data Figure 3.**
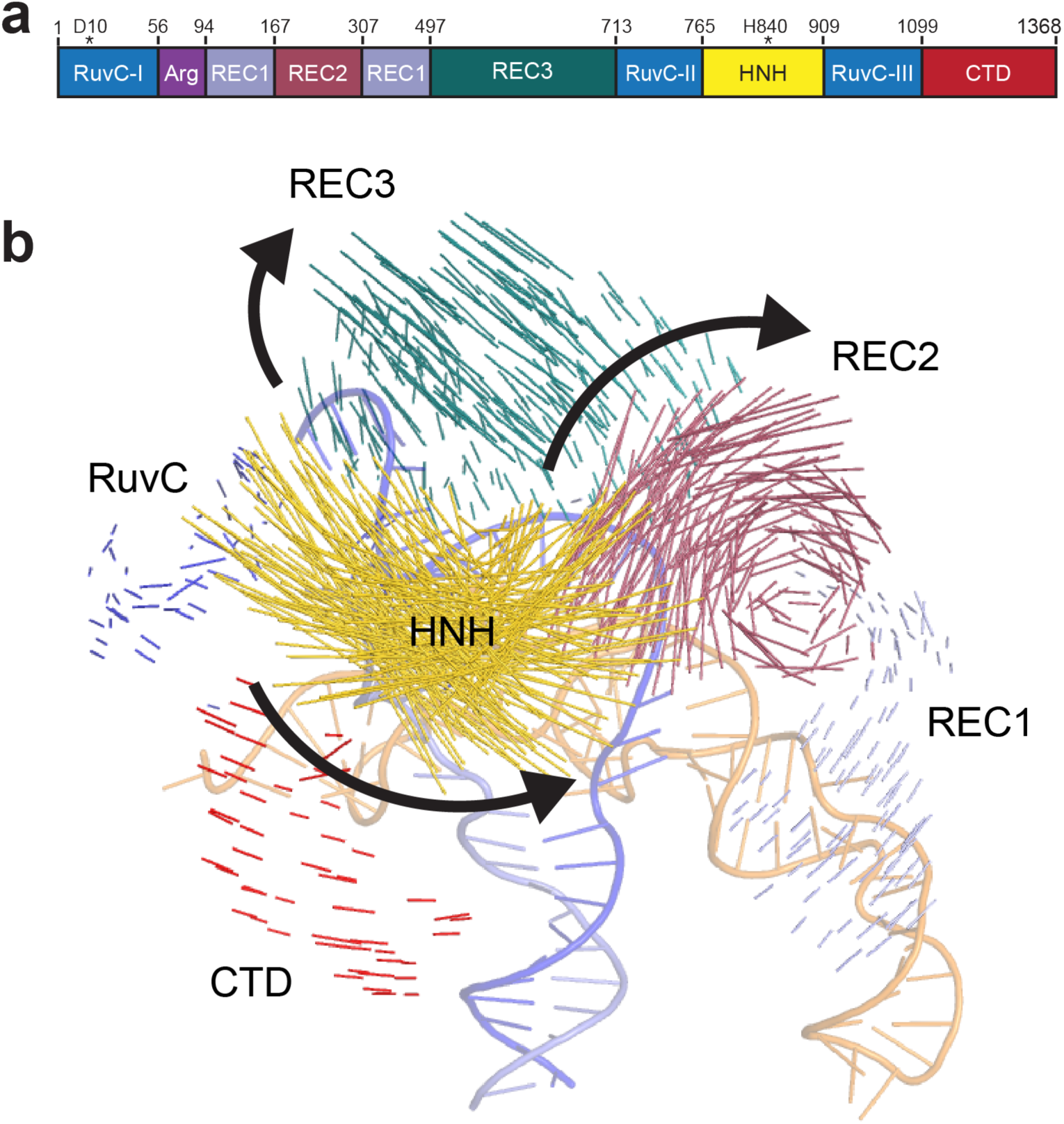
The HNH nuclease, REC2 and REC3 domains undergo substantial conformational changes upon binding to the dsDNA target. **a,** Schematic of SpCas9 domain structure with color coding for separate domains. **b,** Vector map of global SpCas9 conformational changes from the sgRNA- (PDB ID: 4ZT0) to dsDNA-bound structures (PDB ID: 5F9R), domains colored as in panel **a**.

**Extended Data Figure 4.**
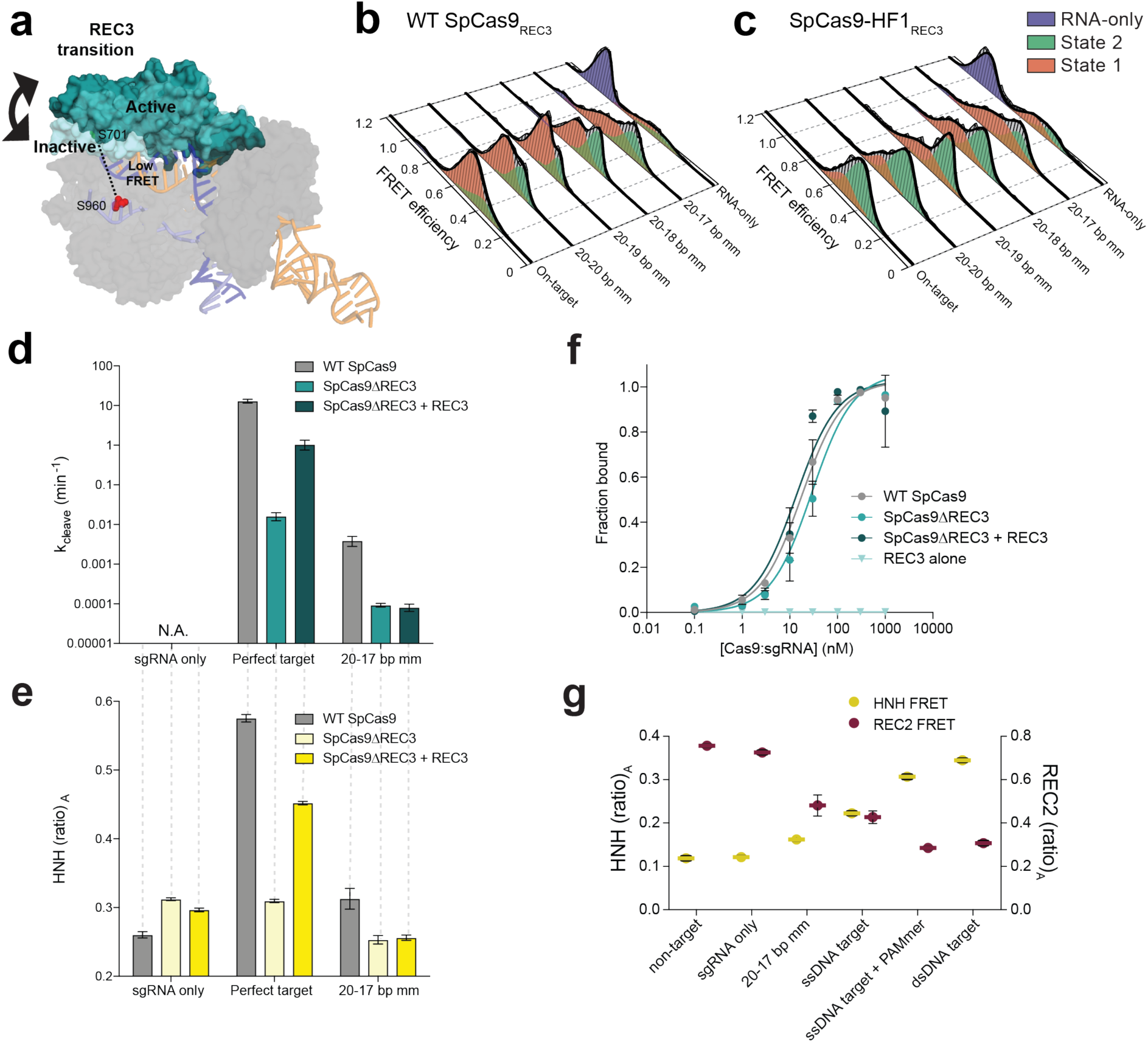
Nucleic acid sensing requires engagement with the REC3 domain and outward rotation of the REC2 domain. **a,** Schematic of SpCas9_REC3_ with FRET dyes at positions S701C and S960C, with HNH domain omitted for clarity. Inactive to active structures represent REC3 in the sgRNA-bound (PDB ID: 4ZT0) to dsDNA-bound (PDB ID: 5F9R) forms, respectively. **b–c,** smFRET histograms measuring HNH conformational activation with black curves representing a fit to multiple Gaussian peaks for **b,** WT SpCas9_REC3_ and **c,** SpCas9-HF1_REC3_ bound to perfect and PAM-distal mismatched targets. The purple peak denotes the sgRNA-only bound state, while the red and green peaks represent two states of REC3 with conformational flexibility upon binding to DNA substrates. **d–e,** REC3 *in vitro* complementation assay with SpCas9ΔREC3 by measuring **d**, cleavage rate constants and **e,** HNH activation with (ratio)_A_ values. **f,** Perfect target DNA binding assay in the presence or absence of the REC3 domain. **g,** (Ratio)_A_ data with SpCas9_REC2_ and SpCas9_HNH_ showing reciprocal FRET states with the indicated substrates. Error bars in **d–g**, s.d.; *n* = 3.

**Extended Data Figure 5.**
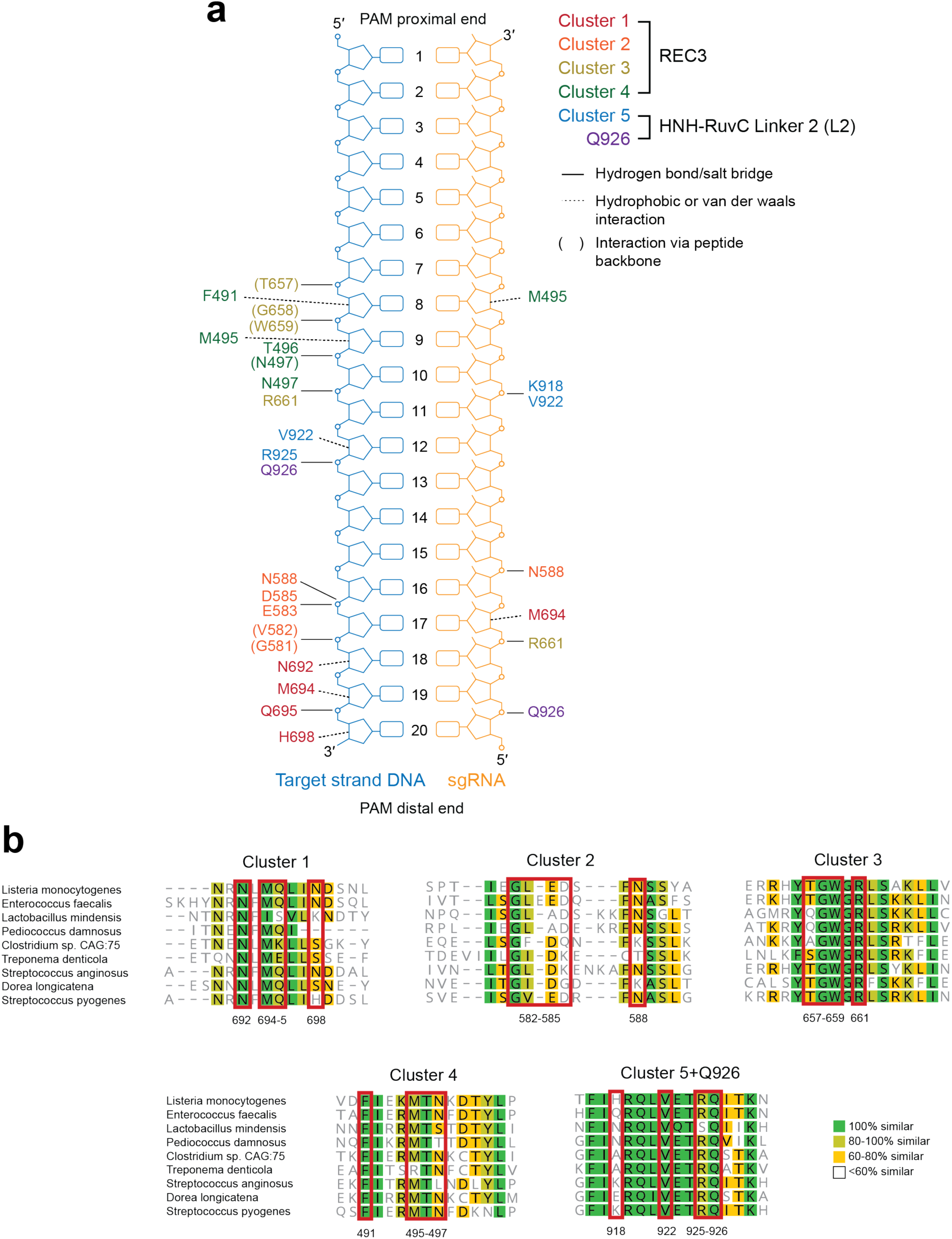
Identification of Cluster variants based on nucleic acid proximity and multiple sequence alignment of residues within Clusters 1-5. **a,** Schematic depicting interactions of WT SpCas9 residues within Clusters 1-5 with the RNA/DNA heteroduplex, based on PDB accession 5F9R (adapted from ref 8). **b,** Alignment of selected Cas9 orthologues using MAFFT and visualized in Geneious 10.0, with red boxes outlining residues mutated to alanine within each cluster variant.

**Extended Data Figure 6.**
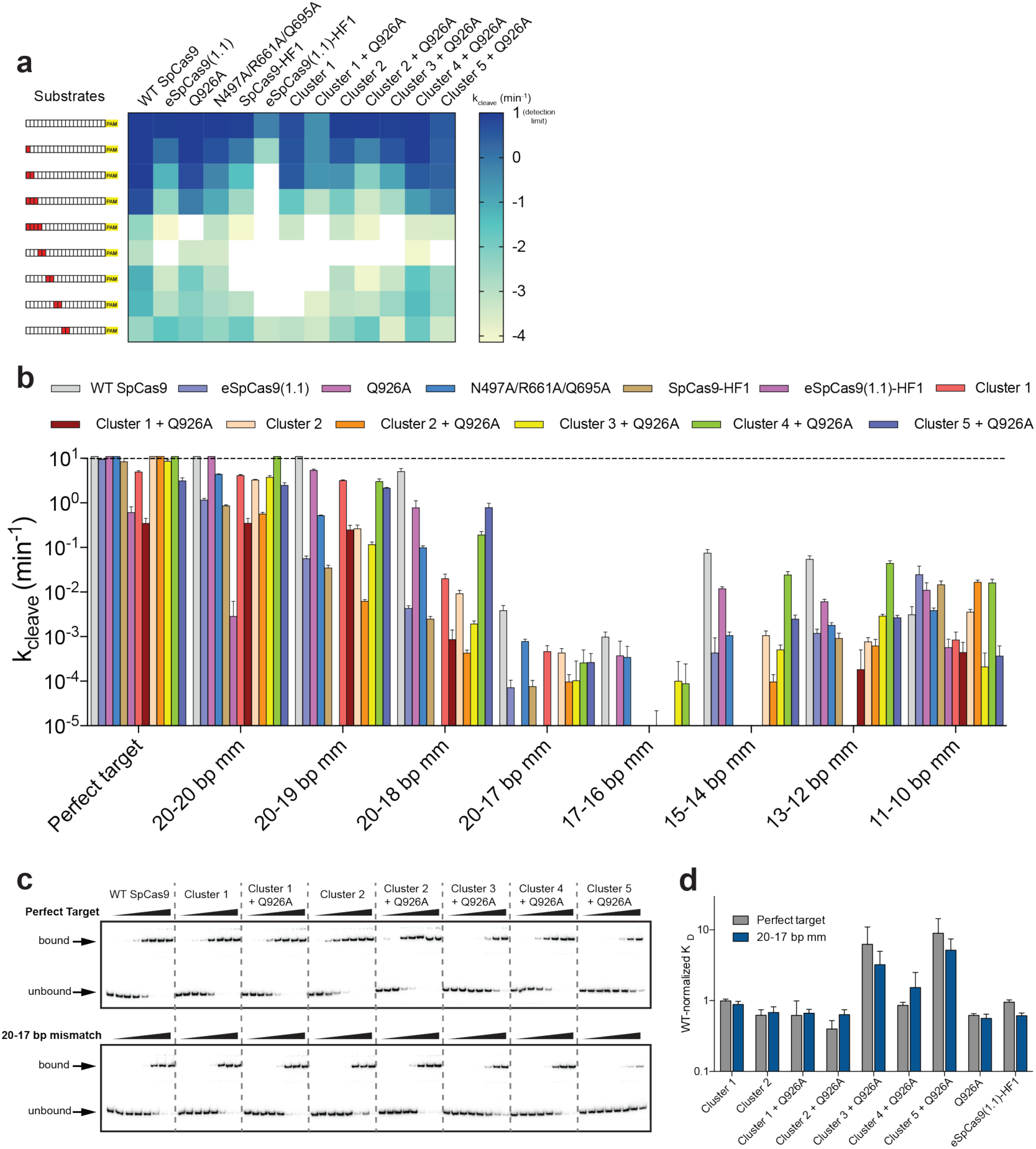
Mutation clusters in the REC3 domain along the RNA/DNA heteroduplex demonstrate localized sensitivity to mismatches along the target sequence. **a- b,** Quantified DNA cleavage rates (detection limit for k_cleave_ set at 10 min^-1^) displayed as a **a,** heatmap and **b,** bar graph. **c-d,** Target DNA binding assay **c,** resolved by native polyacrylamide gel electrophoresis (PAGE) mobility shift assays and **d,** quantification with WT-normalized dissociation constants. Error bars in **b** and **d**, s.d.; *n* = 3.

**Extended Data Figure 7.**
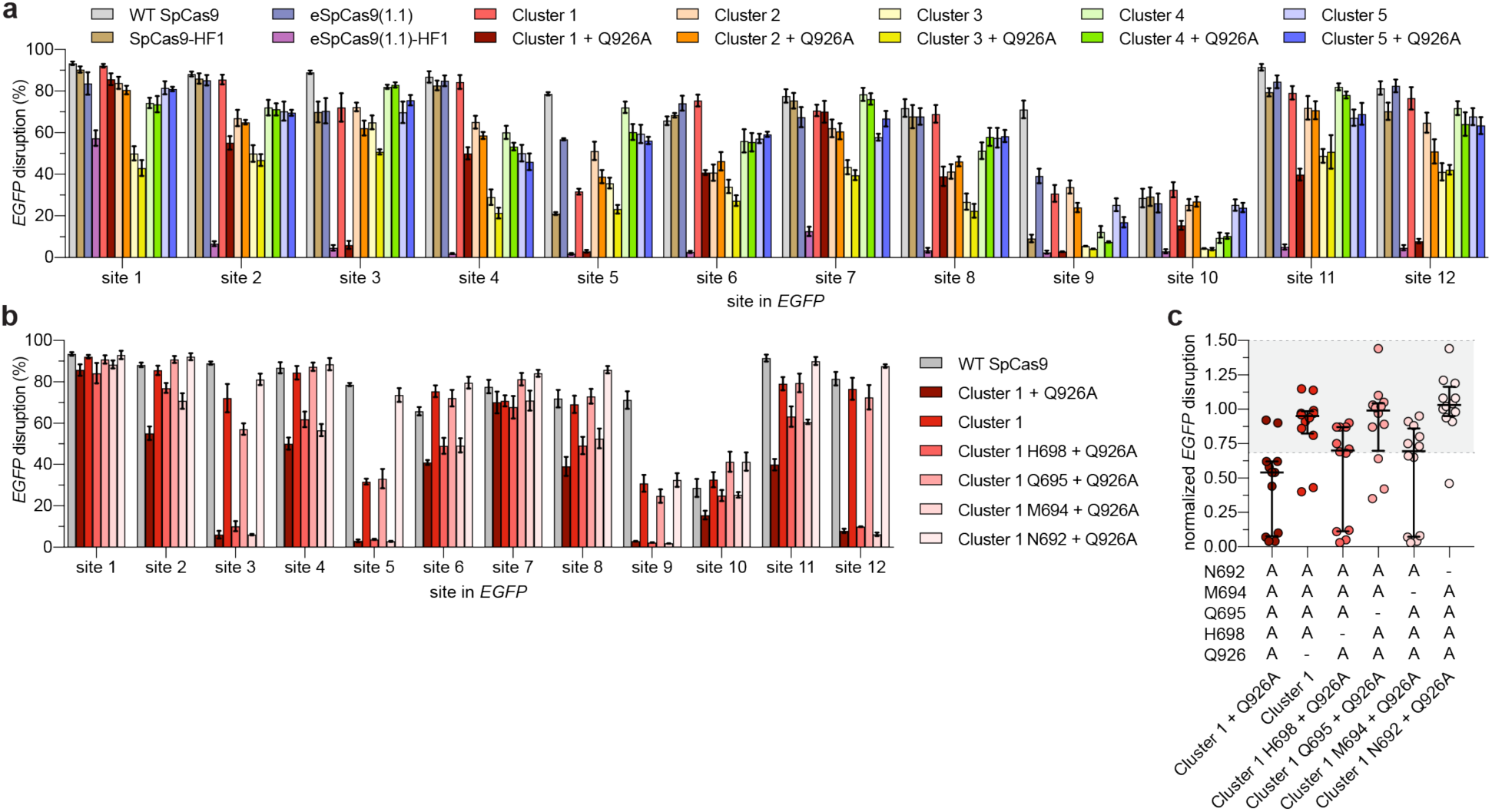
On-target activities of altered specificity variants using a human cell *EGFP* disruption assay. **a,** Summary of *EGFP* disruption activities for SpCas9-HF1, eSpCas9(1.1), eSpCas9(1.1)-HF1 and Cluster variants ± Q926A with mean and s.e.m., where n ≥ 3. **b,** Summary of *EGFP* disruption activities for the series of Cluster 1 variants with each substituted residue restored to the canonical amino acid; mean and s.e.m. where n ≥ 3; WT, Cluster 1, and Cluster 1 A926Q data from **panel a** is re-plotted for comparison. **c,** WT- normalized plot of data in **panel b**; error bars represent median and interquartile range; the interval with > 70% of wild-type activity is highlighted in light grey.

**Extended Data Figure 8.**
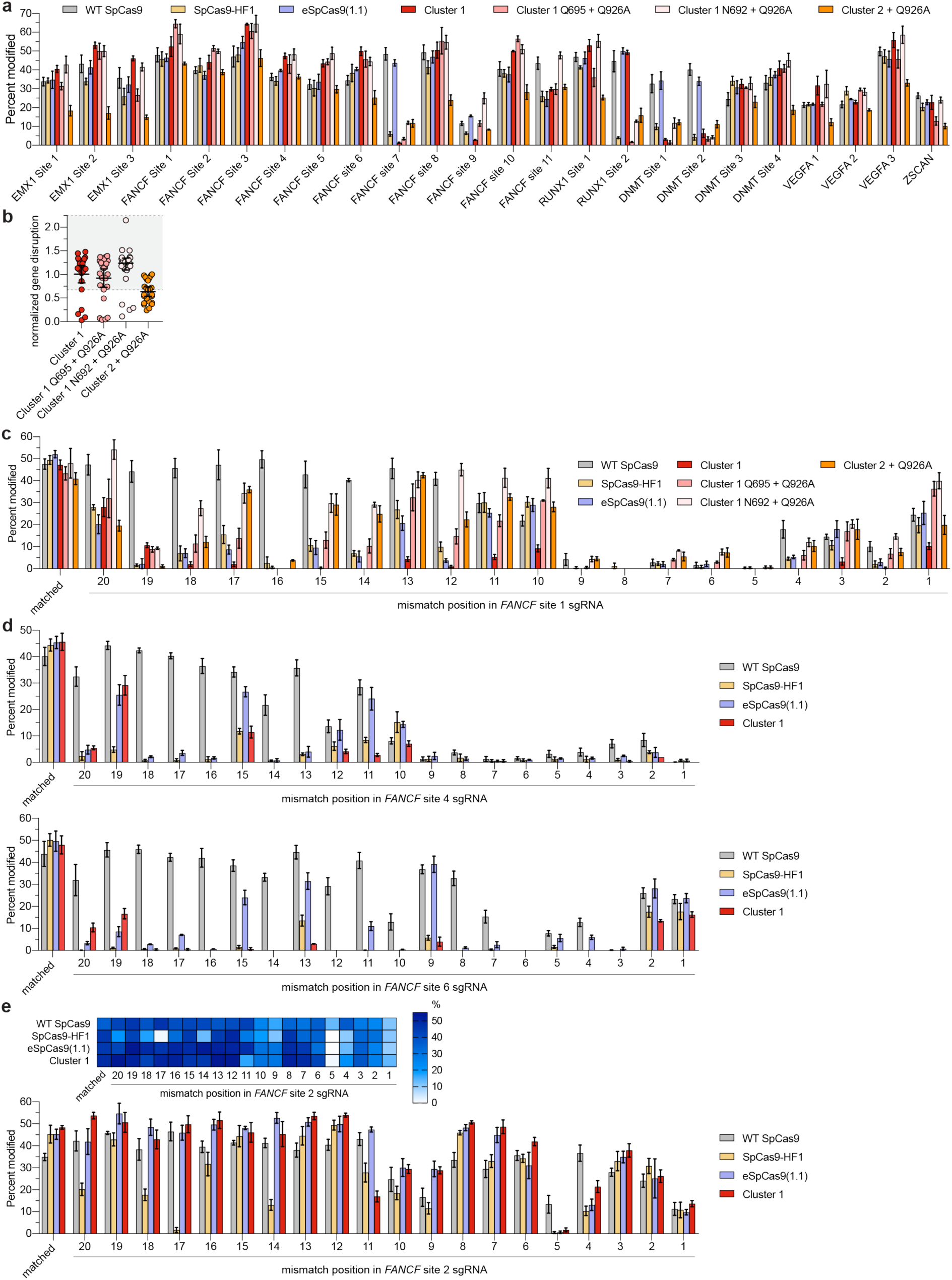
Activities and specificities of high-fidelity SpCas9 variants targeted to endogenous human cell sites. **a,** On-target activities of WT SpCas9, SpCas9-HF1, Cluster 1 and Cluster 2 variants across 24 endogenous human genes, assessed by T7E1 assay. Mean and s.e.m. shown; n ≥ 3. **b,** WT-normalized endogenous gene disruption data from panel **a**, for Cluster 1 and 2 variants. Error bars represent median and interquartile ranges with the > 70% interval of wild-type activity highlighted in light grey; Cluster 1 A926Q data from **Fig. 3b** is replotted for comparison. **c-e,** Summary of single mismatch tolerance of WT SpCas9, SpCas9-HF1, eSpCas9(1.1), and Cluster 1 and Cluster 2 variants on **c,** *FANCF* site 1 **d,** *FANCF* sites 4 and 6, and **e,** *FANCF* site 2. Percent modification assessed by T7E1 assay; mean and s.e.m. shown; n ≥ 3.

**Extended Data Figure 9.**
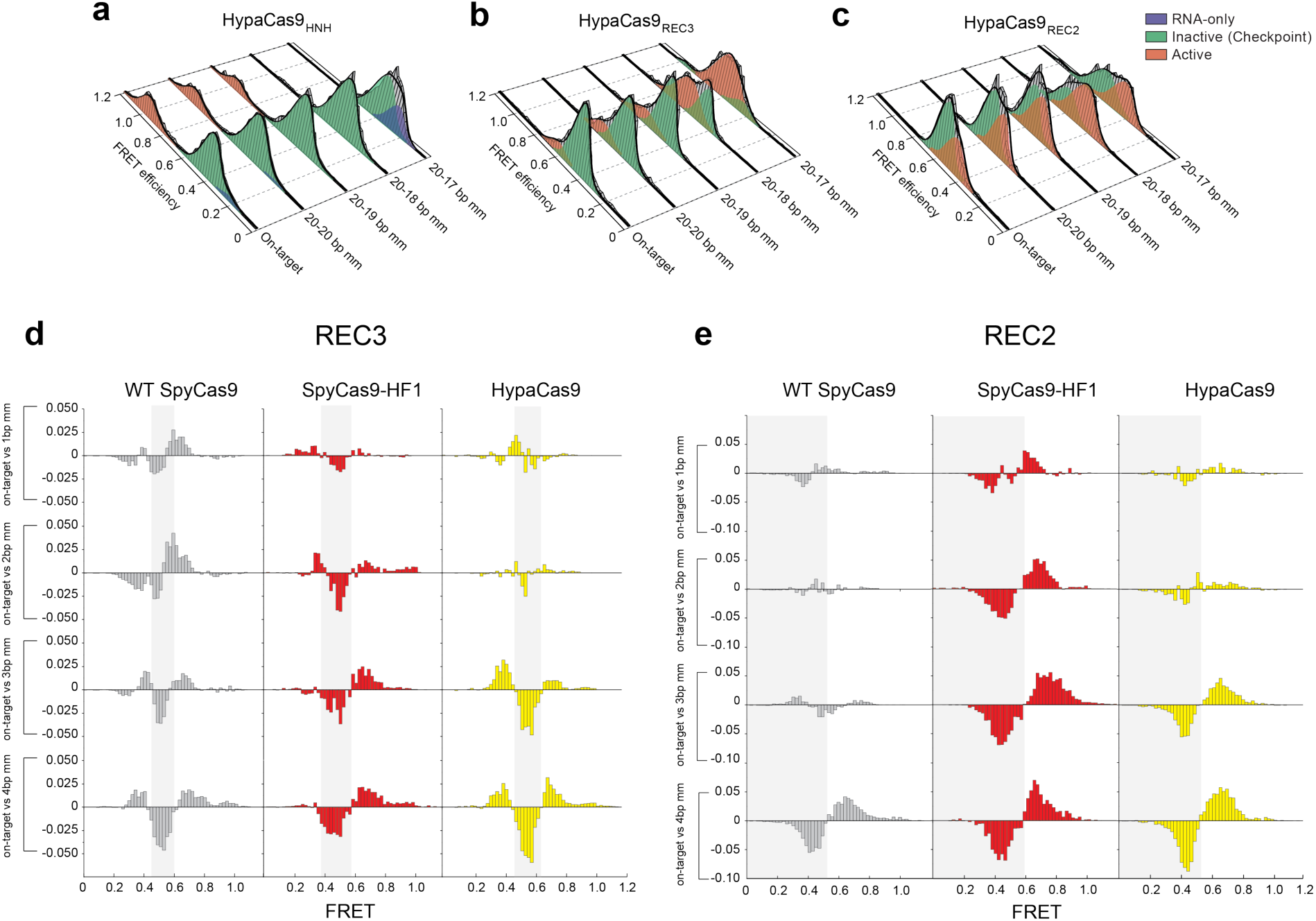
Conformational gating drives targeting accuracy for SpCas9 variants. **a-c, Steady state** smFRET histograms measuring **a,** HNH, **b,** REC2 and **c,** REC3 conformational states for HypaCas9 bound to on-target and PAM-distal mismatched substrates. Black curves represent a fit to multiple Gaussian peaks. **d-e,** Steady state smFRET histograms of Cas9 variants bound to PAM distal mismatched substrates were normalized to and subtracted from that of on-target smFRET histograms. This analysis reveals transitions from one FRET population (negative peak) to another population (positive peak) for **d,** REC3 and **e,** REC2.

**Extended Data Table 1.** Measured distances between residues labelled with Cy3/Cy5 FRET dyes for different substrate-bound Cas9 structures. Residue pairs were designed to report conformational changes of the specified domain (HNH, REC2 or REC3). The distances were measured between Cα atoms of the indicated residues for the associated PDB structures.

|  |  | Inter-residue distance (Å) |  |  |
| --- | --- | --- | --- | --- |
|  |  | HNH | REC2 | REC3 |
| Structure | PDB | (S355-S867) | (E60-D273) | (S701-S960) |
| sgRNA-only | 4ZT0 | 81 | 12.8 | 28.4 |
| sgRNA/TS DNA-bound | 4OO8 | 61 | 35.9 | 45.1 |
| sgRNA/TS DNA + PAMmer-bound | 4UN3 | 59 | 37.2 | 46.5 |
| sgRNA/dsDNA-bound | 5F9R | 30 | 32 | 40.3 |

**Supplementary Table 1.** All enhanced specificity, high-fidelity, cluster and hyper-accurate SpCas9 variants tested in this study. The HNH, REC3 or REC3 subscript designation with an enhanced specificity, high-fidelity or cluster SpCas9 variant denotes combination of residue substitutions with indicated FRET construct.

| Protein description | Substitution/Truncation |  |  |  |  |  |  |
| --- | --- | --- | --- | --- | --- | --- | --- |
|  | 1 | 2 | 3 | 4 | 5 | 6 | 7 |
| Wild-type SpCas9 | - | - | - | - | - | - | - |
| SpCas9<sub>HNH</sub> | C80S | C574S | S355C | S867C | - | - | - |
| SpCas9<sub>REC2</sub> | C80S | C574S | E60C | D273C | - | - | - |
| SpCas9<sub>REC3</sub> | C80S | C574S | S701C | S960C | - | - | - |
| SpCas9ΔREC3 | M1–N497, GGS, V713–D1368 |  |  |  | - | - | - |
| SpCas9ΔREC3<sub>HNH</sub> | M1–N497, GGS, V713–D1368 |  |  |  | + HNH FRET |  | - |
| SpCas9ΔREC3<sub>REC2</sub> | M1–N497, GGS, V713–D1368 |  |  |  | + REC2 FRET |  | - |
| REC3 domain | K506–Q712 |  | - | - | - | - | - |
| SpCas9-Q926A | Q926A | - | - | - | - | - | - |
| SpCas9-N497A/R661A/Q695A | N497A | R661A | Q695A | - | - | - | - |
| SpCas9-N497A/R661A/Q695A<sub>HNH</sub> | N497A | R661A | Q695A | + HNH FRET |  | - | - |
| SpCas9-HF1 | N497A | R661A | Q695A | Q926A | - | - | - |
| SpCas9-HF1<sub>HNH</sub> | N497A | R661A | Q695A | Q926A | + HNH FRET |  | - |
| SpCas9-HF1<sub>REC2</sub> | N497A | R661A | Q695A | Q926A | + REC2 FRET |  | - |
| SpCas9-HF1<sub>REC3</sub> | N497A | R661A | Q695A | Q926A | + REC3 FRET |  | - |
| SpCas9-K855A | K855A | - | - | - | - | - | - |
| SpCas9-K855A<sub>HNH</sub> | K855A | + HNH FRET |  | - | - | - | - |
| eSpCas9(1.1) | K848A | K1003A | R1060A | - | - | - | - |
| eSpCas9(1.1)<sub>HNH</sub> | K848A | K1003A | R1060A | + HNH FRET |  | - | - |
| eSpCas9(1.1)-HF1 | N497A | R661A | Q695A | Q926A | K848A | K1003A | R1060A |
| Cluster 1 (HypaCas9) | N692A | M694A | Q695A | H698A | - | - | - |
| Cluster 1<sub>HNH</sub> | N692A | M694A | Q695A | H698A | Q926A | + HNH FRET |  |
| Cluster 1<sub>REC2</sub> | N692A | M694A | Q695A | H698A | Q926A | + REC2 FRET |  |
| Cluster 1<sub>REC3</sub> | N692A | M694A | Q695A | H698A | Q926A | + REC3 FRET |  |
| Cluster 1 + Q926A | N692A | M694A | Q695A | H698A | Q926A | - | - |
| Cluster 1 H698 + Q926A | N692A | M694A | Q695A | - | Q926A | - | - |
| Cluster 1 Q695 + Q926A | N692A | M694A | - | H698A | Q926A | - | - |
| Cluster 1 M694 + Q926A | N692A | - | Q695A | H698A | Q926A | - | - |
| Cluster 1 N692 + Q926A | - | M694A | Q695A | H698A | Q926A | - | - |
| Cluster 2 | G582A | V583A | E584A | D585A | N588A | - | - |
| Cluster 2 + Q926A | G582A | V583A | E584A | D585A | N588A | Q926A | - |
| Cluster 3 | T657A | G658A | W659A | R661A | - | - | - |
| Cluster 3 + Q926A | T657A | G658A | W659A | R661A | Q926A | - | - |
| Cluster 4 | F491A | M495A | T496A | N497A | - | - | - |
| Cluster 4 + Q926A | F491A | M495A | T496A | N497A | Q926A | - | - |
| Cluster 5 | K918A | V922A | R925A | - | - | - | - |
| Cluster 5 + Q926A | K918A | V922A | R925A | Q926A | - | - | - |

**Supplementary Table 2.**
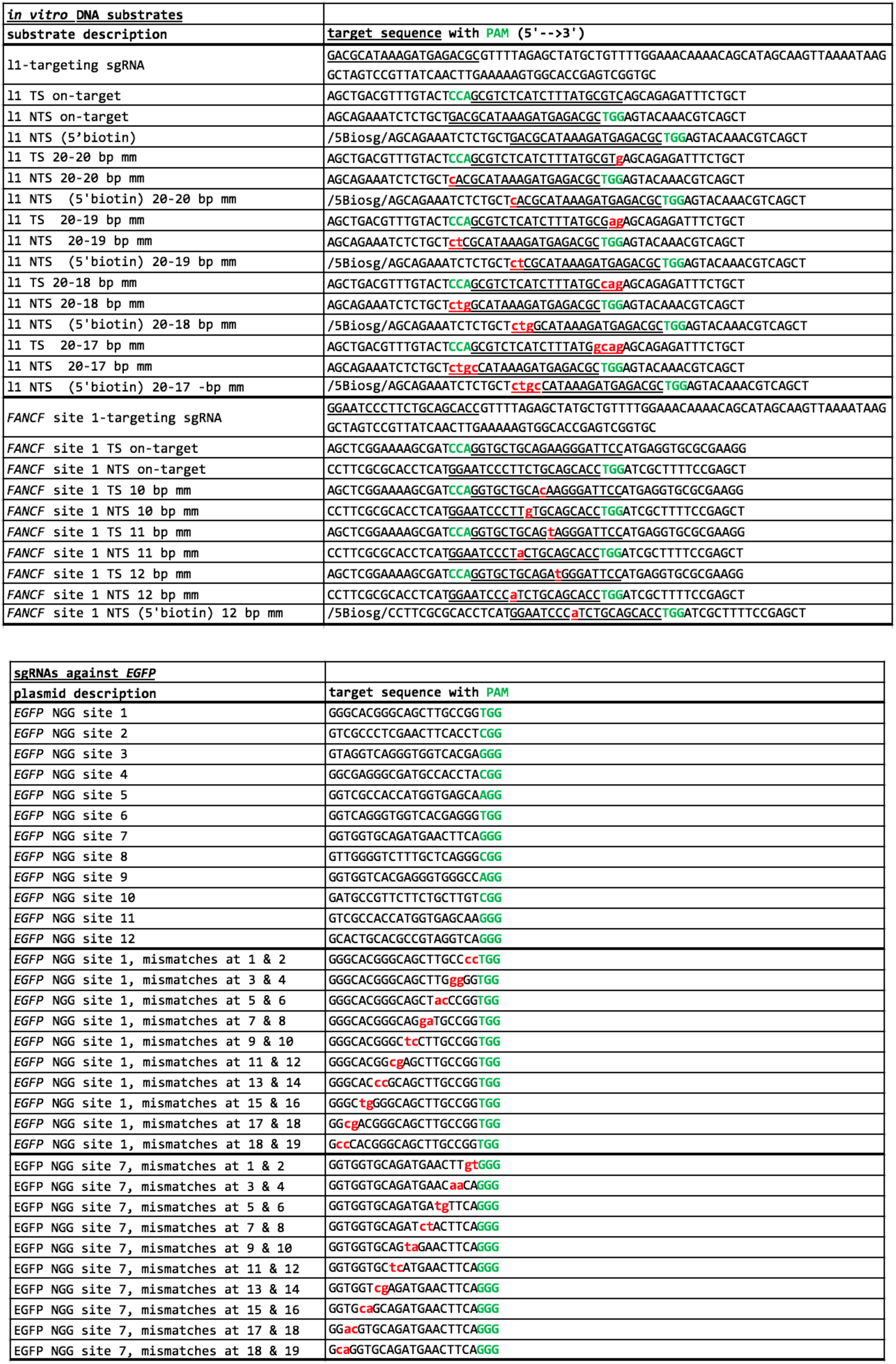
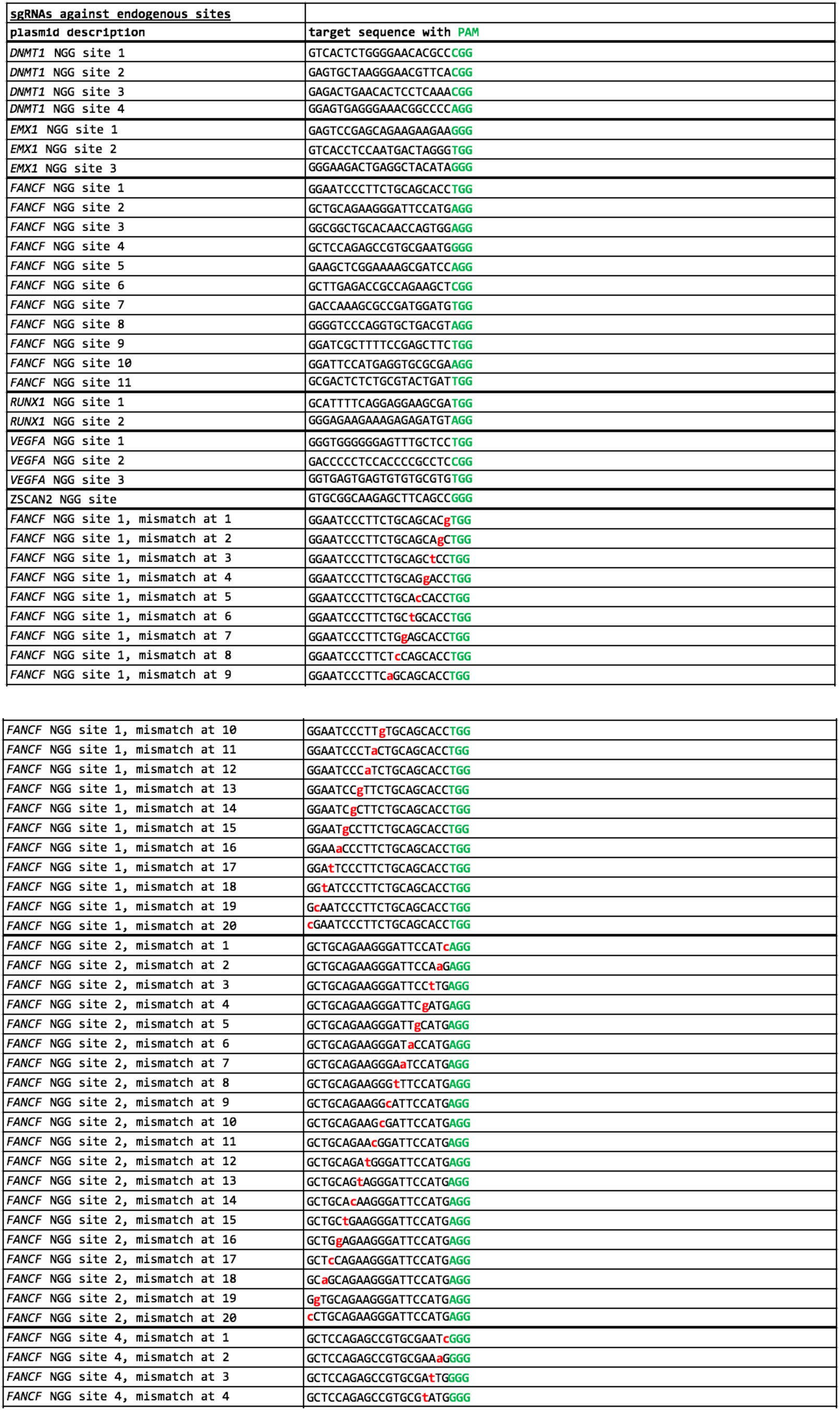
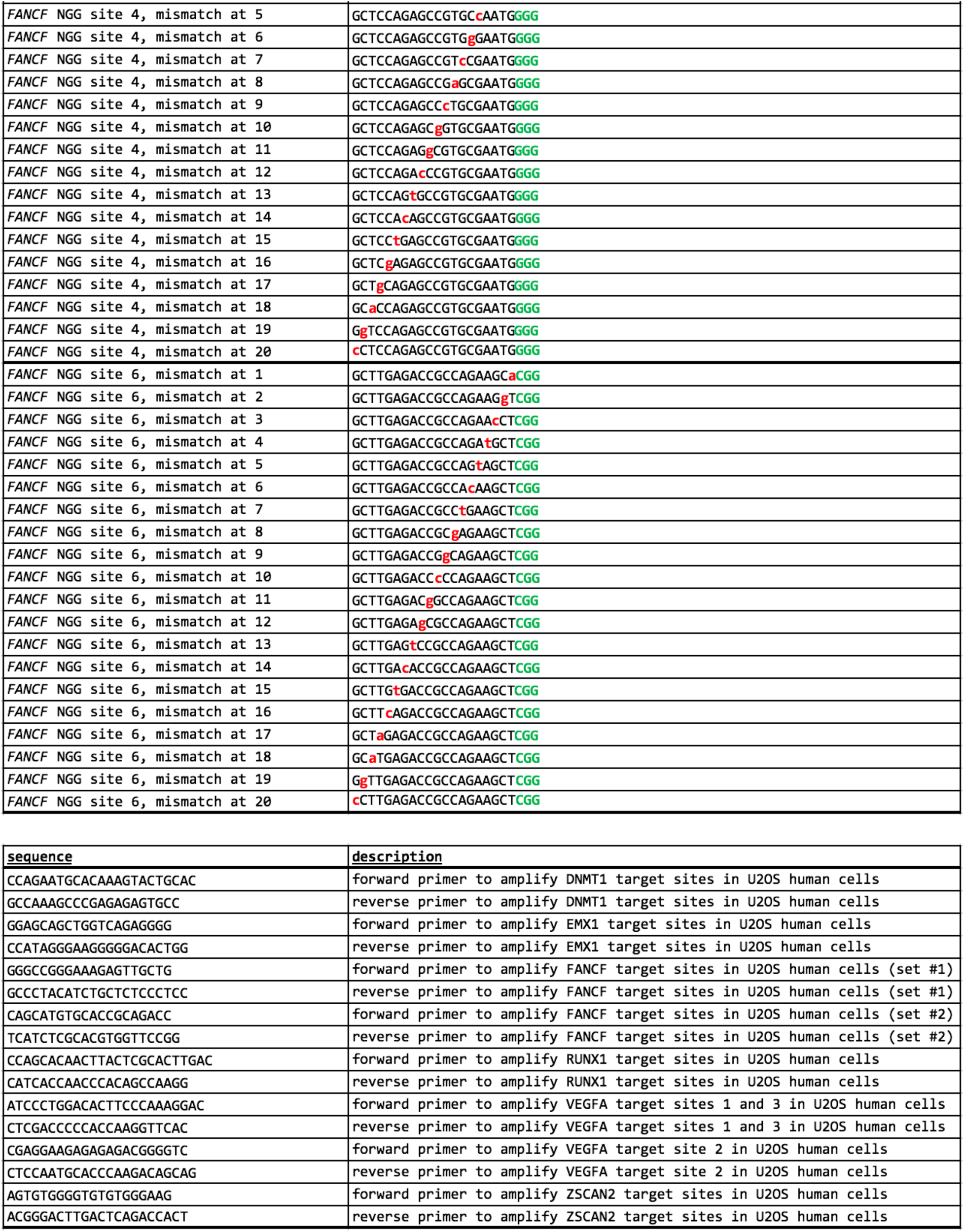
Sequences of all nucleic acids used in the study.

